# A transient hypothalamic CRH signal coordinates social investigation in mice

**DOI:** 10.64898/2026.05.14.725080

**Authors:** Ibukun Akinrinade, Meenakshi Pardasani, Toni-Lee Sterley, Tamás Füzesi, Jaideep S. Bains

## Abstract

Successful navigation of the social environment depends on the ability to rapidly evaluate conspecifics and adjust behaviour accordingly. Encounters with unfamiliar individuals are particularly challenging because they introduce uncertainty about how the interactions may unfold. Such social encounters can lead to beneficial or detrimental outcomes, making rapid assessment critical for survival; yet the neural mechanisms underlying this process remain poorly understood. Corticotropin-releasing hormone synthesizing neurons in the paraventricular nucleus of the hypothalamus (CRH^PVN^), traditionally viewed as the canonical endocrine controllers of the stress response, have been implicated in processing behaviourally relevant environmental and social information, including the investigation of stressed conspecifics. However, their role in rapidly evaluating conspecifics during social encounters remains unclear. Here, using fibre photometry and optogenetics during ethologically relevant social encounters, we show that CRH^PVN^ neurons exhibit a rapid, transient increase in activity that precedes investigative behaviour. This signal scales with social context, is elevated during exposure to unfamiliar relative to familiar conspecifics, persists across novel environments, and generalizes across unfamiliar strains. In contrast, interactions with juvenile conspecifics and inanimate objects do not elicit this differential activity, indicating that CRH^PVN^ recruitment is not driven solely by novelty. Critically, temporally specific inhibition of CRH^PVN^ neurons during this brief activity window produces lasting changes in behaviour beyond the period of inhibition, including the reduction of appropriate social investigation, indicating that this transient activity is necessary for gating subsequent social behaviour. Together, these findings identify CRH^PVN^ neurons as a rapid integrative node linking hypothalamic function to social evaluation.

## INTRODUCTION

Animals continuously encounter diverse social and environmental cues and must rapidly evaluate them to guide appropriate behaviour. In humans and other social species, encounters with unfamiliar individuals require rapid appraisal to determine the likely outcome of the interaction, a process that is critical for guiding behaviour and ensuring safety^1,2^. Social interactions can be highly beneficial, supporting development, reproduction, and emotional resilience, yet they can also introduce challenges during competition for limited resources, excessive aggression and the establishment of dominance hierarchies^3–8^. Importantly, these social encounters are often ambiguous, requiring rapid evaluation under conditions of uncertainty^9–11^. Failure to appropriately assess social cues can lead to maladaptive outcomes, either through delayed responses to adverse situations or through unnecessary behavioural and physiological costs associated with overestimation of risk^12,13^. Despite the importance of these processes, the neural mechanisms that support rapid social assessment remain poorly understood.

Emerging evidence implicates corticotropin-releasing hormone neurons in the paraventricular nucleus of the hypothalamus (CRH^PVN^), classically known for orchestrating endocrine stress responses, in processing behaviourally relevant information beyond canonical stress regulation^14–18^. Previous work has shown that CRH^PVN^ neurons are required for social investigation of stressed conspecifics, a behaviour that facilitates the transmission of affective state between individuals, and that these neurons are engaged by social stimuli^19^. However, whether CRH^PVN^ neurons contribute to the rapid evaluation of conspecifics to guide appropriate behavioral responses during social encounters remains unclear.

Here, using fibre photometry and optogenetics in an ethologically relevant social interaction paradigm, we demonstrate that CRH^PVN^ neurons exhibit rapid, transient increases in activity that are critical for subsequent investigative behaviours. This activity pattern suggests a role in gating behavioural responses that enable the assessment of conspecifics. Together, our findings identify CRH^PVN^ neurons as a key component of the neural circuitry underlying rapid social appraisal, linking endocrine regulation to real-time behavioural decision-making.

## RESULTS

### Increased CRH^PVN^ neuron activity during social appraisal of an unfamiliar conspecific

Social interactions vary in predictability. Unfamiliar conspecifics introduce social uncertainty demanding rapid evaluation to guide appropriate behavioral responses^9,20^. To investigate how CRH^PVN^ neural activity contributes to this process, we used transgenic mice expressing GCaMP6f selectively in CRH neurons^17^ (Fig. 1A). We implanted an optical fibre above the PVN to perform fibre photometry recordings of calcium-dependent fluorescence as a proxy for neuronal population activity. Using a classical resident-intruder test, the neural activity was specifically recorded from the resident mouse during social interaction. Following baseline recording of the resident mouse during a solitary period, a familiar same-sex cage mate or an unfamiliar same-sex, same-strain intruder was introduced into the resident’s home cage (Fig. 1B). Approach toward a familiar intruder elicited a transient increase in CRH^PVN^ activity, which returned to baseline within seconds and remained around baseline during the remainder of the 3-min interaction period (Fig. 1C). This activity emerged within seconds of intruder introduction across conditions, indicating rapid engagement of CRH^PVN^ neurons during the initial stages of social evaluation (Fig. 1D-F). Fine-grained temporal analysis of each condition revealed that although the introduction of both familiar and unfamiliar intruders produced rapid CRH^PVN^ activity, the introduction of an unfamiliar intruder triggered a significantly greater and more sustained early response (Fig. 1G). The peak amplitude, peak area under the curve (AUC) and time to baseline of the averaged CRH^PVN^ Ca^2+^ responses in the resident mouse were significantly larger when exposed to unfamiliar intruders than to familiar intruders (Fig. 1H-J). Likewise, this differential response occurred in both male and female resident animals (Fig. S1A-B). Together, these findings demonstrate that CRH^PVN^ neurons are recruited during social appraisal in a manner that scales with familiarity status, supporting a role for these neurons as rapid detectors of ambiguous social information.

**Figure 1.**
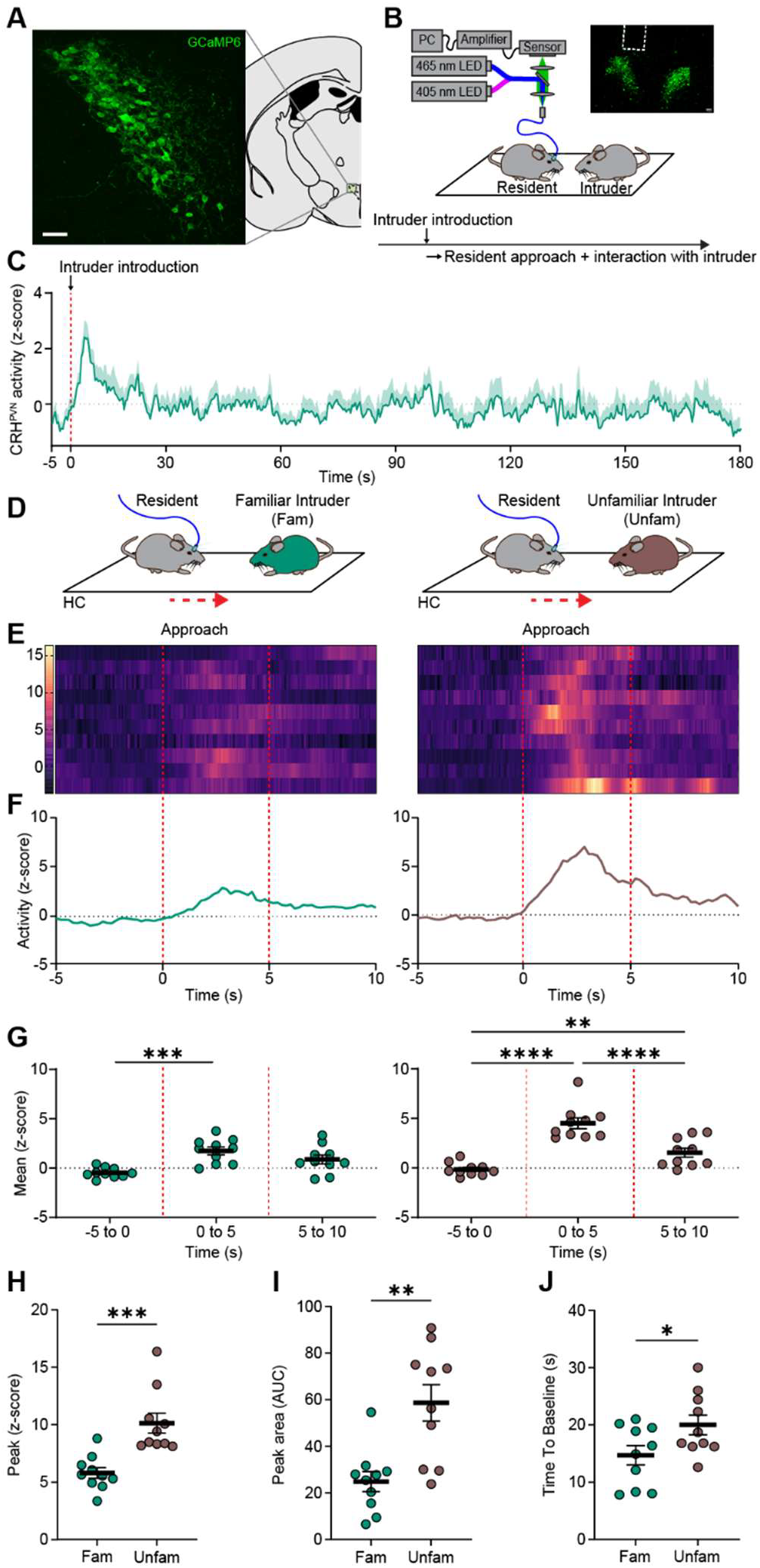
Increase in CRH^PVN^ activity during conspecific intruder appraisal. **A**. Coronal hemi-section illustrating the paraventricular nucleus (PVN) of the hypothalamus (right) and corresponding confocal image (left) demonstrating GCaMP6f expression in CRH-expressing PVN neurons (scale bar = 50 µm). **B**. Schematic of single-fiber photometry recordings from CRH^PVN^ neurons during social investigation. Confocal image (right) shows the ferrule position above the PVN (scale bar = 100 µm), with the social interaction paradigm illustrated below. **C**. Mean Ca_2+_ activity traces from CRH^PVN^ neurons during social assessment of a familiar conspecific (n = 10). **D**. Schematic depicting a resident mouse in the home cage (HC) approaching familiar (green) and unfamiliar (brown) intruders, with approach direction indicated by a dashed red arrow. **E**. Approach-aligned heatmaps of CRH^PVN^ Ca_2+_ responses, with each row representing an individual animal and the corresponding mean z-scored activity trace shown below (**F**). **G**. Mean z-scored CRH^PVN^ Ca_2+_ activity across 5-second time windows (−5 to 0, 0 to 5, 5 to 10 s), with time 0 indicating resident approach towards intruder (n = 10). Left: familiar intruder (repeated-measures one-way ANOVA, F(1.653,14.87) = 11.55, p = 0.001; Bonferroni’s multiple comparisons: −5 to 0 vs 0 to 5, p < 0.0001, t = 6.316; −5 to 0 vs 5 to 10, p=0.0626, t=2.795; 0 to 5 vs 5 to 10, p = 0.4272, t = 1.606). Right: unfamiliar intruder (repeated-measures one-way ANOVA, F(1.445,13.00) = 58.81, p < 0.0001; Bonferroni’s multiple comparisons: −5 to 0 vs 0 to 5, p < 0.0001, t = 8.511; −5 to 0 vs 5 to 10, p = 0.008, t = 4.116; 0 to 5 vs 5 to 10, p < 0.0001, t = 9.405). **H-J**. Comparison of the first post-approach calcium transients between familiar and unfamiliar intruder exposures (n = 10 mice per group). Peak amplitude: p = 0.0004, t = 4.393 (**H**); peak area under the curve (AUC): p = 0.0013, t = 3.803 (**I**); time to baseline: p = 0.04, t = 2.200 (**J**; unpaired t-tests).

### Differential CRH^PVN^ neuron activity persists in novel contexts and across conspecific strains

To determine whether the heightened CRH^PVN^ responses to unfamiliar conspecifics reflect social uncertainty rather than context-specific factors, we examined neuronal activity in a neutral non-territorial environment. To this end, we placed mice in a novel cage with fresh bedding and performed fibre photometry recording during exposure to both familiar and unfamiliar intruders (Fig. 2A).

**Figure 2.**
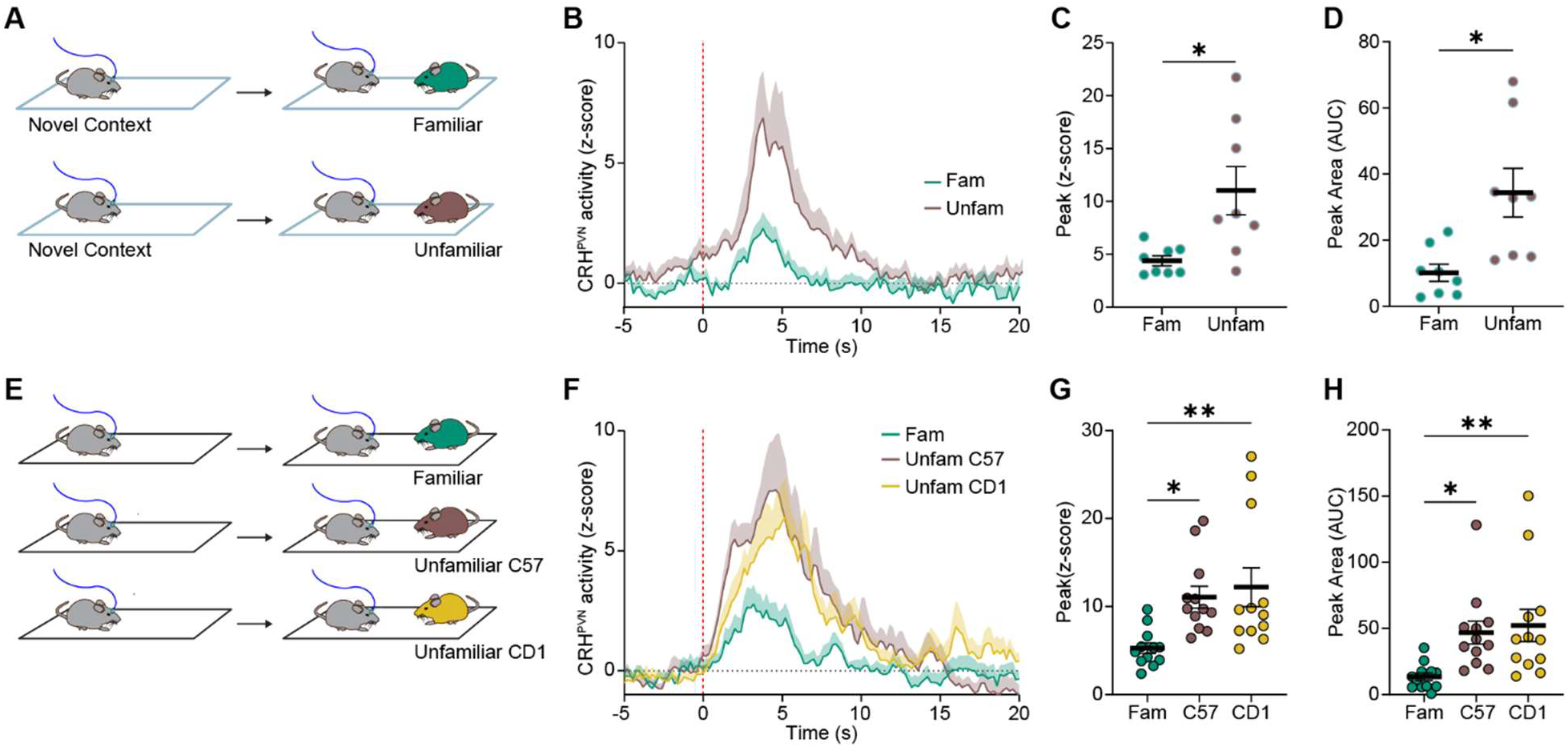
Differential CRH^PVN^ neuron activity persists in novel context and across conspecific strains. **A**. Schematic illustrating a resident mouse exposed to familiar (top) and unfamiliar intruders (bottom) in a novel context. **B**. Group-averaged z-scored CRH^PVN^ Ca_2+_ responses aligned to approach, with shaded area representing s.e.m. **C**. Comparison of individual peak z-scores for the first Ca_2+_ transient following approach (p = 0.0274, t = 2.776; n = 8 mice per group; unpaired t-test). **D**. Comparison of peak area under the curve (AUC) for the first Ca_2+_ transient following approach (n = 8 mice per group; p = 0.0118, t = 3.376; unpaired t-test). **E**. Schematic depicting resident interactions with familiar and unfamiliar same-strain (C57) intruders and a non–same-strain (CD1) intruder in the home cage. **F**. Group-averaged z-scored CRH^PVN^ Ca_2+_ responses aligned to approach, with shaded area representing s.e.m. **G**. Individual peak z-scores of the first calcium transient following approach (ordinary one-way ANOVA, F(2,33) = 2.154, p = 0.0056; Tukey’s multiple comparisons: Fam vs Unfam-C57 p = 0.0271, q = 3.849; Fam vs Unfam-CD1 p = 0.0073, q = 4.597; Unfam-C57 vs Unfam-CD1 p = 0.8575, q = 0.7487; n = 12 mice per group). **H**. Individual peak area under the curve (AUC) of the first calcium transient following approach (ordinary one-way ANOVA, F(2,33) = 5.743, p = 0.0072; Tukey’s multiple comparisons: Fam vs Unfam-C57 p = 0.0289, q = 3.809; Fam vs Unfam-CD1 p = 0.01, q = 4.424; Unfam-C57 vs Unfam-CD1 p = 0.9014, q = 0.6149; n = 12 mice per group).

Consistent with observations made in the home cage, approach toward unfamiliar intruders in a novel context elicited a significantly greater increase in CRH^PVN^ activity compared to familiar interactions (Fig. 2B, S2A,B). The peak amplitude and AUC of CRH^PVN^ responses were significantly higher during interactions with unfamiliar compared to familiar mice (Fig. 2C,D), indicating that enhanced CRH^PVN^ responses persist independently of environmental familiarity. We next asked whether this differential activity generalizes across conspecific types. In the home cage, interactions with either unfamiliar same-strain (C57) or non– same-strain (CD1) intruders elicited significantly greater CRH^PVN^ activity compared to familiar mice (Fig. 2E,F, S3A,B). Analysis of the first calcium transient revealed increased responses to both unfamiliar C57 and CD1 intruders relative to familiar partners, but no difference between unfamiliar groups (Fig. 2G,H). Together, these findings demonstrate that CRH^PVN^ neuron activity robustly scales with conspecific unfamiliarity across contexts and strain differences, supporting a role for these neurons in detecting social uncertainty during the initial stages of social appraisal.

### Differential CRH^PVN^ neuron activity is absent during exposure to unfamiliar juvenile intruders and object

To determine whether CRH^PVN^ activity reflects unfamiliarity or is selectively engaged under social conditions requiring evaluation, we compared responses across interactions with familiar adult, unfamiliar adult, and unfamiliar juvenile conspecifics. While approach toward unfamiliar adult intruders elicited an increase in CRH^PVN^ calcium activity relative to familiar adult interactions, responses to unfamiliar juvenile intruders were not elevated and were comparable to those observed during familiar adult encounters (Fig. 3A–D, S4A-B). These findings indicate that enhanced CRH^PVN^ activity is not a general feature of novelty but is selectively recruited during interactions with unfamiliar adult conspecifics.

**Figure 3.**
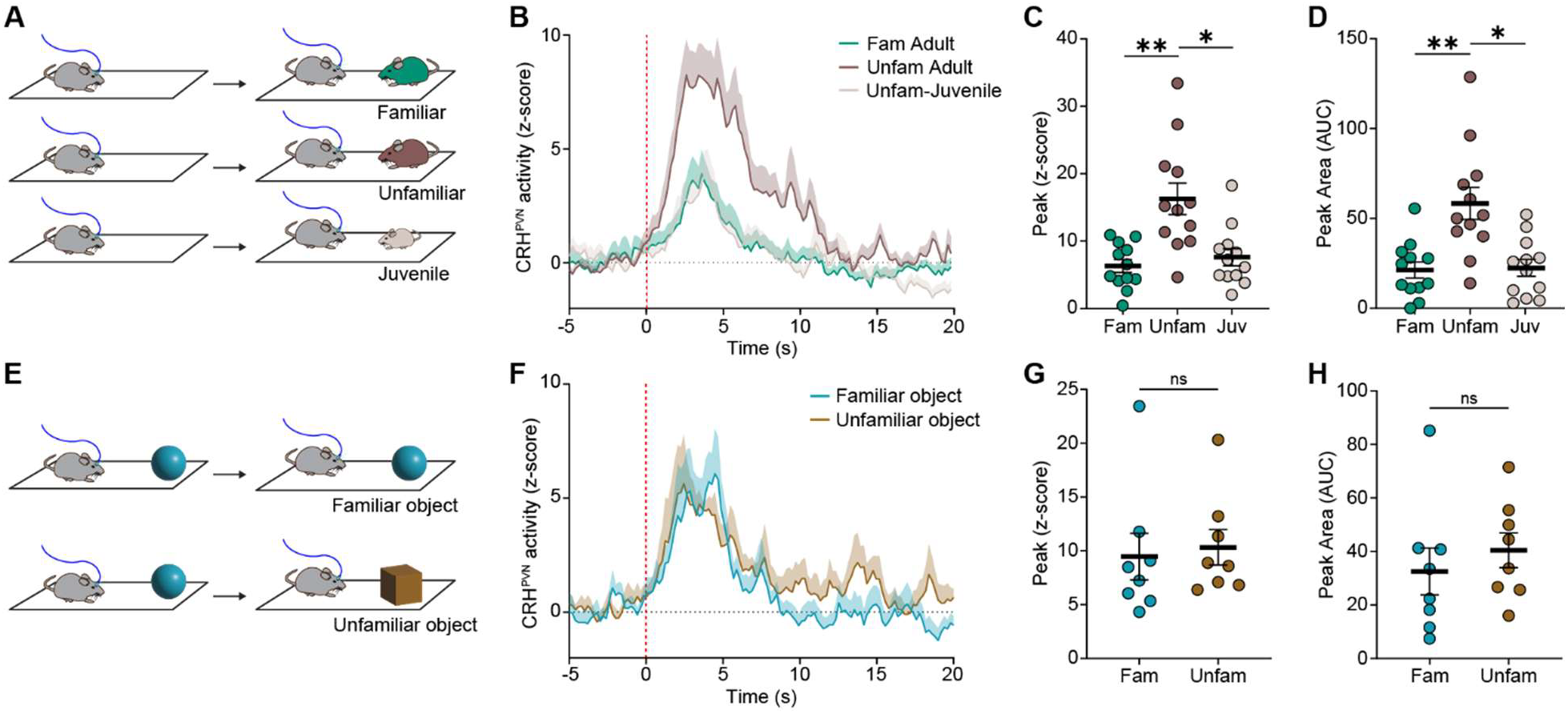
CRH^PVN^ activity is similar during exposure to unfamiliar juvenile intruders and objects. **A**. Schematic depicting exposure of resident to familiar, unfamiliar, and juvenile intruders in home cage. **B**. Group average z-score of CRH^PVN^ calcium responses aligned at approach, shade represents s.e.m. **C**. Individual peak z-scores of the first peak after approach (Ordinary 1-way ANOVA, F (2,35) = 10.02, p = 0.0059, Tukey’s multiple comparison test, Fam-Adult vs. Unfam-Adult p = 0.0095, q=5.185, Unfam-Adult vs. Unfam-Juvenile p = 0.0378, q=4.051, and Fam-Adult vs. Unfam-Juvenile, p = 0.4343, q=1.811, *n* = 12 mice per group). **D**. Individual peak area under the curve for the calcium activity for the first peak after approach (Ordinary 1-way ANOVA, F (2,35) = 11.87, p = 0.0041, Tukey’s multiple comparison test, Fam-Adult vs. Unfam-Adult p = 0.0083, q=5.298, Unfam-Adult vs. Unfam-Juvenile p = 0.0178, q=4.669, and Fam-Adult vs. Unfam-Juvenile, p = 0.9124, q=0.5792, *n* = 12 mice per group). **E**. Schematic depicting resident interactions with familiar and unfamiliar objects in homecage. **F**. Group average z-score of CRH^PVN^ calcium responses aligned at approach, shade represents s.e.m. **G**. Individual peak z-scores of the first peak after approach (*p* = 0.4284, t = 0.8404, *n* = 12 mice per group, unpaired, *t*-test). **H**. Individual peak area under the curve for the calcium activity for the first peak after approach (*p* = 0.4256, t = 0.8458, *n* = 12 mice per group, unpaired, *t*-test).

We next assessed whether this activity pattern extends to non-social stimuli. Following a 3-day pre-exposure to the familiar object, resident mice were exposed to either a familiar or a novel object^21,22^.

During the approach to the familiar and unfamiliar objects, CRH^PVN^ activity did not differ between conditions (Fig. 3E–H). Together, these results demonstrate that while CRH^PVN^ neurons are broadly responsive to novelty or unfamiliar stimuli, they are differentially selectively engaged in social contexts that require evaluation of unfamiliar adult conspecifics, supporting a role in context-dependent social appraisal.

### Increased social investigation during exposure to unfamiliar conspecifics

The selective increase in CRH^PVN^ activity towards unfamiliar intruders raises the question of whether this neural signal carries behavioral consequences. To address this, we first sought to determine how conspecific familiarity shapes behavioural engagement during social encounters, we systematically quantified distinct behavioural classes during interactions with familiar and unfamiliar conspecifics. Raster plots revealed the temporal structure of discrete behavioural events, including sniffing, rearing, digging, and grooming, across the interaction period (Fig. 4A). These behaviours provide a comprehensive assessment of distinct social and non-social components of the interaction. Sniffing reflects direct social investigation and active information sampling of conspecifics^23–25^, whereas rearing, digging, and grooming provide measures of general exploration, arousal, and self-directed behaviour^26–28^. This distinction allowed us to isolate behaviours specifically related to social assessment. Across conditions, sniffing behaviour was initiated immediately upon exposure to the intruders and was predominantly directed from the resident toward the intruder, reflecting active investigation of the social partner (Fig. 4Bi–ii). Likewise, the total duration of resident-to-intruder sniffing was increased during interactions with unfamiliar conspecifics compared to familiar conspecifics (Fig. 4Biii). This enhancement in directed investigation indicates that exposure to unfamiliar conspecifics triggers increased sampling of socially relevant cues, consistent with a greater need for social assessment. In contrast, non-social behaviours, including digging, rearing, and grooming, were not significantly altered between familiar and unfamiliar conditions (Fig. S5A-C), and the investigation of novel objects was reduced compared to familiar objects (Fig. S5D), indicating that the observed differences are specific to social investigation.

**Figure 4.**
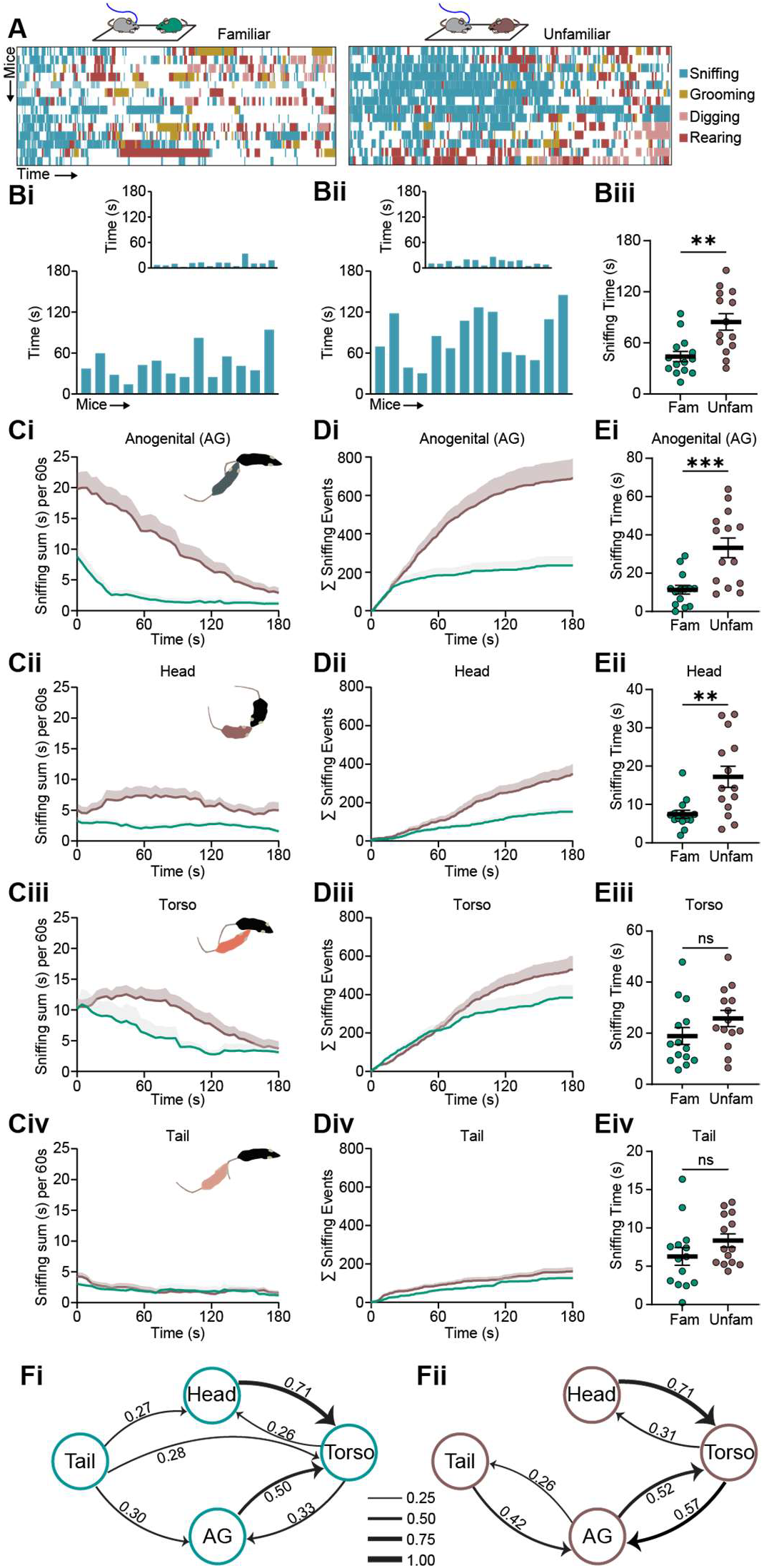
Increased social investigation during exposure to unfamiliar intruder. **A**. Raster plots illustrating the temporal distribution of behavioural events following exposure to familiar (left) and unfamiliar (right) intruders. Each row corresponds to an individual animal, with event markers denoting discrete behavioural events over time. **B**. Vertical bars showing individual sniffing durations for resident-to-intruder (bottom) and intruder-to-resident (top) interactions under familiar (**Bi**) and unfamiliar (**Bii**) conditions. Total sniffing duration during unfamiliar investigations is higher compared to familiar investigations (**Biii**) (*p* = 0.0015, t = 3.550, *n* = 14 mice per group, unpaired, *t*-test). **C**. Temporal dynamics of sniffing behavior during social investigation, quantified as sliding sums of anogenital (**Ci**), head (**Cii**), torso (**Ciii**), and tail (**Civ**) sniffing time, averaged in 60-second bins with a 3-second sliding window. Cumulative sum of sniffing events during social investigations, showing total time spent in anogenital (**Di**), head (**Dii**), torso (**Diii**), and tail (**Div**) sniffing. Differential sniffing behavior during familiar and unfamiliar intruder investigation, quantified across anogenital (**Ei**), head (**Eii**), torso (**Eiii**), and tail (**Eiv**) regions. (Anogenital: *p* = 0.0006, t = 3.896, *n* = 14 mice per group, unpaired *t*-test; Head: *p* = 0.0028, t = 3.299, *n* = 14 mice per group, unpaired *t*-test; Torso: *p* = 0.1459, t = 1.499, *n* = 14 mice per group, unpaired *t*-test; Tail: *p* = 0.1604, t = 1.445, *n* = 14 mice per group, unpaired *t*-test). (**F**). Transition probability plots depicting the dynamics of switching between investigative behaviours during interaction with familiar (**i**) and unfamiliar (**ii**) intruders during social investigation. Nodes correspond to behavioural states, and directed edges are weighted by transition probabilities.

To further resolve the structure of this investigation, we decomposed sniffing into anatomically defined subtypes corresponding to anogenital, head, torso, and tail regions. This breakdown was motivated by the functional relevance of these regions, as different targets provide distinct social information, including identity, reproductive status, threat level, and recent social experience^19,29–35^. Temporal analysis revealed that sniffing behaviour is dynamically distributed over the interaction period, with distinct patterns emerging across sniffing subtypes (Fig. 4Ci-iv). Cumulative measures showed that increases in sniffing during exposure to unfamiliar intruders were not uniform but selectively enriched for specific investigative behaviours (Fig. 4Di-iv). Specifically, the investigation of unfamiliar intruders was characterized by increased anogenital and head-directed sniffing. In contrast, torso and tail sniffing did not differ substantially between conditions, indicating that unfamiliarity selectively enhanced behaviours most relevant to social evaluation rather than broadly increasing all forms of contact (Fig. 4Ei-iv). Permutation-based comparison of behavioural transition matrices revealed a significant difference in overall transition architecture between familiar and unfamiliar conditions (Frobenius norm = 0.375, p = 0.0044). The observed statistic exceeded the 95th percentile of the null distribution, indicating that the behavioural structure differed beyond chance levels.

Inspection of the difference matrix (unfamiliar − familiar) showed that this effect was driven by structured changes in specific transitions rather than uniform shifts across the network. The most prominent increase was observed in transitions from torso to anogenital sniffing (+0.25), with additional increases in transitions from tail to anogenital sniffing (+0.11). In contrast, transitions from torso to tail (−0.16) and torso self-transitions (−0.14) were reduced. Edgewise statistical testing with false discovery rate correction identified a subset of significant transitions, all originating from the torso state. These included an increased probability of transitioning from torso to anogenital sniffing and decreased probabilities of transitions from torso to tail and torso itself (Fig. 4Fi-ii, S5E-G). Together, these results indicate that unfamiliar animals exhibit a reorganization of behavioural transitions characterized by a preferential routing of behavioural sequences toward anogenital sniffing, along with suppression of alternative transitions. Our results therefore highlight anogenital sniffing as a key behavior for social assessment.

### CRH^PVN^ activity is necessary for targeted anogenital investigation during social assessment

Our behavioral analysis revealed enhanced investigative sniffing directed towards unfamiliar intruders. We next asked if the rapid, transient increase in the CRH^PVN^ activity neurons during social approach causally drives the behavioral response. To test this, we expressed Archaerhodopsin 3.0 virus in CRH neurons to optogenetically inhibit CRH^PVN^ neuron activity (Fig. 5A). Targeted viral expression and fibre implantation enabled selective silencing of CRH^PVN^ neurons, which was validated by ex vivo slice recordings demonstrating reliable light-driven inhibition of neuronal firing^36^ (Fig. 5B). During the experiment, optogenetic inhibition of the resident mouse was done during the initial phase of interaction (20s), corresponding to the period when CRH^PVN^ activity rapidly increased upon intruder exposure. Behavioural quantification was performed to capture the entire 3 min interaction period. We first quantified the overall social investigation by measuring the total sniffing behaviour in Arch-inhibited and mCherry expressing controls. Under control conditions, interactions with unfamiliar conspecifics elicited greater overall sniffing than interactions with familiar conspecifics. However, optogenetic inhibition of CRH^PVN^ neurons selectively reduced total sniffing during exposure to unfamiliar intruders, without altering investigation of familiar intruders (Fig. 5C).

**Figure 5.**
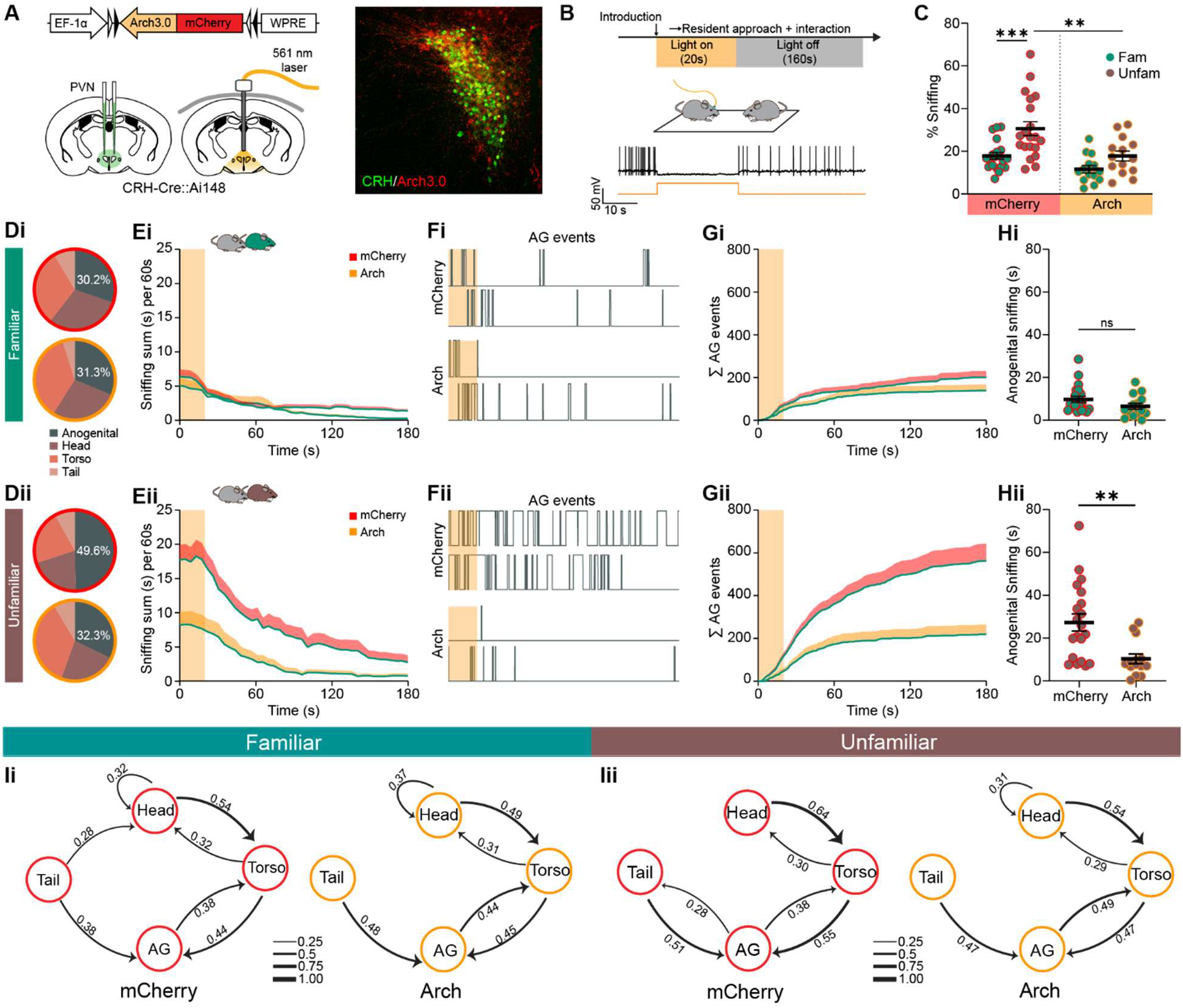
Increase in CRH^PVN^ activity is essential for targeted anogenital investigation. A. Top: Viral construct for optogenetic inhibition experiment, bottom: Strategy of virus injection and ferrule implantation for optogenetic inhibition of CRH^PVN^ neurons, right: Confocal image depicting Cre-dependent localization of aav2/9-EF1a-DIO-Arch3.0 mcherry virus in the CRH^PVN^ neurons (Scale bar = 50 μm). **B**. Experimental timeline depicting photo-inhibition of CRH^PVN^ neurons in the resident mouse during the first 20s of the social investigation of an intruder; and representative whole-cell recording from a CRH^PVN^ neuron in a brain slice, demonstrating spike suppression in response to 561-nm light. **C**. Percentage of total sniffing behavior during social investigation, comparing control (mCherry) and optogenetically inhibited (Arch) residents during investigation of familiar and unfamiliar intruders (n=14-20 per group, 1-way ANOVA, F(4,68) = 11.18, p < 0.0001, Holm-Sidak’s multiple comparison test, Fam-mCherry vs Unfam-mCherry p= 0.0007, t=4.003; Fam-mCherry vs Fam-Arch p = 0.1492, t=1.794, Unfam-mCherry vs. Unfam-Arch p = 0.0016, t=3.505; Fam-Arch vs Unfam-Arch p=0.1492, t= 1.634). **D**. Pie charts showing the distribution of sniffing behavior in control (mCherry) and optogenetically inhibited (Arch) residents. (**i**) Similar distribution during investigation of a familiar intruder. (**ii**) Altered distribution during unfamiliar intruder investigation. **E**. Temporal profile of anogenital sniffing during social investigation, shown as mean sniffing time averaged in 60-second bins with a 3-second sliding window, comparing mCherry controls and Arch-inhibited groups during (**i**) familiar and (**ii**) unfamiliar intruder investigation. **F**. Example raster plots of anogenital sniffing events for (**i**) familiar and (**ii**) unfamiliar intruders (top: mCherry-control; bottom: Arch-inhibited). **G**. Cumulative anogenital sniffing dynamics during investigation of (**i**) familiar and (**ii**) unfamiliar intruders, shown as group-averaged duration in mCherry controls and Arch-inhibited animals. **H**. Comparison of anogenital sniffing time between mCherry controls and Arch-inhibited residents, shown as mean ± s.e.m., during investigation of (**i**) familiar intruders (p = 0.1353, t = 1.532) and (**ii**) unfamiliar intruders (p = 0.0023, t = 3.307; unpaired t-tests). **I**. Transition matrices illustrating the probability of switching between investigative behaviours during interaction with familiar (**i**) and unfamiliar (**ii**) intruders, comparing mCherry controls (left) and Arch-inhibited residents (right).

To further resolve how CRH^PVN^ activity shapes social assessment, we decomposed total sniffing into distinct investigative subtypes, including anogenital, head, torso, and tail-directed sniffing. This breakdown is important because these behaviours reflect different components of social investigation^29,31,33,34^. In control animals, unfamiliar interactions were characterized by a distinct distribution of these behaviours, indicative of targeted investigation. In contrast, inhibition of CRH^PVN^ neurons disrupted this organization, leading to an altered distribution of sniffing behaviours specifically during unfamiliar encounters, while leaving familiar interaction patterns intact (Fig. 5Di-ii, S6A-F).

We next examined the temporal dynamics of investigation, focusing on anogenital sniffing as a key index of sustained social evaluation^19,33^ (Fig. S5F). In control animals, unfamiliar conspecifics elicited prolonged, temporally structured anogenital investigation. Inhibition of CRH^PVN^ neurons markedly reduced this sustained engagement, as reflected in both the temporal profile and the sum of cumulative anogenital sniffing events (Fig. 5E–Hi-iii). Analysis of behavioural transitions demonstrated that while CRH^PVN^ inhibition reduced time spent on anogenital sniffing, no corresponding changes in the sequence of social behaviours were observed. Across both familiar and unfamiliar conditions, we found no differences between mCherry and Arch animals in either global transition structure or individual behavioral transitions. Observed variations were consistent with chance-level fluctuations as determined by permutation testing (Fig. 5Ii-ii). This suggests that inhibition of CRH^PVN^ neurons alters the overall duration of specific social behaviors without reorganizing the flow of behavioral transitions.

## DISCUSSION

Social interactions require animals to rapidly evaluate conspecifics and adjust behaviour accordingly. Here, we identify CRH^PVN^ neurons as a rapid, transient signal engaged during this process, with activity elevated during encounters with unfamiliar, relative to familiar conspecifics. This response is consistent across environments and strains, but absent in juveniles or non-social objects. Importantly, though transient, this signal is necessary for targeted social investigation. Together, these findings position CRH^PVN^ neurons as a fast, integrative node for assessing conspecifics when interaction outcomes are uncertain.

Neuroimaging studies across humans and rodents have implicated the hypothalamic nuclei as central mediators of socioemotional integration^37^. For example, reproductive and mating-related behaviors engage the medial and ventral hypothalamic nuclei, while predation and defensive evasion recruit the lateral hypothalamic to periaqueductal gray circuitry^38^. These elaborate behaviors, however, are downstream consequences of a more fundamental step: the rapid appraisal of social encounters and the rapid selection of a behavioral response strategy. At this initial decision point, the CRH^PVN^ neurons are well-positioned to play a pivotal role in integrating sensory inputs with internal states, projecting to specialized socially-related hypothalamic nuclei and regulating the neuroendocrine axis, thereby coordinating adaptive responses to environmental challenges^39–42^. Our findings extend this framework by positioning CRH^PVN^ neurons not solely as stress effectors, but as early-stage evaluators of social information. This mechanism may confer an adaptive advantage by enabling individuals the anticipation of potential challenges and the selection of appropriate responses, thereby linking sensory processing to neuroendocrine and behavioral outputs^43–45^.

The behavioural significance of this rapid CRH^PVN^ neural signal is evident in social investigation. Investigative behaviours, particularly anogenital sniffing, are central to how rodents evaluate conspecifics and extract socially relevant cues^46–49^. Our optogenetic inhibition experiments reveal that transient silencing of CRH^PVN^ activity during the early activity window produces lasting changes in investigative behavior that extend beyond the period of inhibition, indicating that this signal gates subsequent behavioural responses. Specifically, inhibition of CRH^PVN^ neuron activity reduced the time animals spent engaged in anogenital investigation, without altering the overall structure of behavioural transitions. This dissociation highlights a primary role for CRH^PVN^ neurons in regulating the intensity and persistence of specific behaviours rather than their sequential organization^50^. Consistent with this interpretation, CRH^PVN^ neurons exert rapid central effects via their projections, in addition to their slower endocrine functions, thereby recruiting downstream circuits that maintain behavioural responses across timescales^51–53^. Given that these neurons also govern the HPA axis^39,54^, it will be important to determine whether their influence on persistent social behaviour arises from synaptic signalling, neuroendocrine modulation, or the coordinated interaction of both systems^55^.

The novel social appraisal functionality of CRH^PVN^ neurons, in addition to their established role in social transmission of stress, defensive flight-fight responses, and anxiety-related behaviors is consistent with the broader mechanistic principles by which PVN operates^14,15,17,19^. Molecularly defined PVN populations form combinatorial assemblies rather than discrete labeled-line circuits^56^, which facilitates the generation of complex behaviors. Within this PVN framework, the CRH neurons likely function not as a single-output relay but as a part of a distributed combinatorial computation that simultaneously coordinates endocrine, behavioral, and autonomic arms of survival responses. How this survival circuit machinery therefore integrates information from the nodes of the social decision-making network remains an important open question. Future work aimed at cell-level encoding, unravelling how distinct CRH^PVN^ neuronal populations govern different aspects of socio-survival processing, will be essential for deepening our mechanistic understanding of social appraisal.

Finally, our study opens new avenues for exploring how CRH^PVN^ neurons contribute to different aspects of social behavior, such as approach, avoidance, and consummatory interactions. More broadly, these neurons may extend the predictive, feed-forward logic seen in homeostatic regulation^57–59^ into the social domain, linking the detection of social uncertainty to the preparation of appropriate internal states. Because disrupted CRH signalling and altered salience processing are key features of stress-related disorders, understanding this mechanism could help guide strategies to restore balanced stress responses and flexible social behaviour, particularly under conditions of social uncertainty.

## METHODS

### Animals

All animal protocols were approved by the University of Calgary Animal Care and Use Committee (AC21-0067). Mice that were the offspring of *Crh-IRES-Cre* (B6(Cg)-*Crh*^*tm1(cre)Zjh*^; stock no: 012704, Jackson Laboratories) and A*i14* (*B6*.*Cg-Gt(ROSA)26Sortm14(CAG-TdTomato)Hze/J*; stock number 007914, Jackson Laboratories) mice crossed with *Ai148 (Ai148(TIT2L-GC6f-ICL-tTA2)-D*; stock number 030328, Jackson Laboratories) were utilized for all fibre photometry and optogenetics with behavioural tracking experiments. Mice were group-housed on a 12:12 h light/dark schedule (lights on at 07:00 hours) with *ad libitum* access to food and water. CD-1 mice (Charles River Laboratory) were used for behavioural alternative strain intruder experiments. Mice were 6–8 weeks old at the time of surgery and virus injection.

### Viruses

AAV carrying Arch3.0-mCherry (LotAAV1081, AAV2/9-EF1a-DIO-Arch3.0-mCherry; Canadian Neurophotonics Platform Viral Vector Core Facility (RRID: SCR_016477) or mCherry (LotAAV1088, AAV2/9-EF1a-DIO-mCherry; Canadian Neurophotonics Platform Viral Vector Core Facility (RRID: SCR_016477)) was used for optogenetic manipulations.

### Stereotaxic injection and optical fibre implantation

Mice were maintained under isoflurane anesthesia in the stereotaxic apparatus. Mono fibre optic cannula with 400 µm diameter (Doric Lenses, MFC_400/430/0.48_5.5mm_MF2.5_FLT) was implanted dorsal to the PVN for fibre photometry (AP, −0.7 mm; L, −0.2 mm from the dura; DV, −4.85 mm from the dura) and for optogenetics (for Arch3.0: AP: −0.7 mm; L: 0.0 mm from the dura; DV: −4.5 mm) following tracking with a custom-made needle (AP: −0.7 mm; L: −0.2 mm from the dura; DV: −4.2 mm) and bilateral injections (AP: −0.7 mm; L: 0.2 mm from the dura; DV: −4.5 mm) fixed to the skull with METABOND® and dental cement. Mice were given 2 weeks to recover prior to the start of the experiment. Experiments started after an additional two weeks of recovery and handling.

### Histology

To verify GCaMP expression and ferrule location, following the experiments, mice were euthanized using isoflurane overdose and a subset of mice were transcardially perfused with phosphate-buffered saline (PBS), followed by 4% paraformaldehyde (PFA) in phosphate buffer (PB, 4 °C). Brains were placed in PFA 24 h followed by 20% sucrose PB. 50 µM coronal brain sections were obtained via cryostat. The other subset of mice was cut into 250 μm sections in ACSF, then fixed in 4% PFA, 1 M PBS, and 30% sucrose for 10 minutes each. Slide-mounted and coverslipped sections were imaged using a confocal microscope (Leica SP8). Images were prepared using ImageJ.

### Slice preparation and electrophysiology

Four to eight weeks post AAV-injection, mice were anesthetized via isoflurane inhalation and decapitated. Rapidly dissected brains were immersed in slicing solution (0–4 °C, 95% O2/5% CO2 saturated) containing (in mM): 87 NaCl, 2.5 KCl, 0.5 CaCl2, 7 MgCl2, 25 NaHCO3, 25 D-glucose, 1.25 NaH2PO4, 75 sucrose. A vibratome (Leica) was used to prepare coronal hypothalamic slices (250 μm thickness), which were transferred for 1+ hours before recording in artificial cerebrospinal fluid (32 °C, 95% O2/5% CO2 saturated) containing (in mM): 126 NaCl, 2.5 KCl, 26 NaHCO3, 2.5 CaCl2, 1.5 MgCl2, 1.25 NaH2PO4, 10 glucose. Recordings were performed in artificial cerebrospinal fluid (1 ml min−1 perfusion) at 30–32 °C. PVN neurons were identified using differential interference contrast and epifluorescence optics (UVICO, Rapp Optoelectronics) and a camera (AxioCam MRm) on an upright microscope (Zeiss). Borosilicate electrodes (3–5 mΩ tip) were backfilled with recording solution composed of (in mM) 108 K-gluconate, 2 MgCl2, 8 Na-gluconate, 8 KCl, 1 K2-EGTA, 4 K2-ATP, 0.3 Na3-GTP, 10 mM HEPES, 10 mg ml−1 biocytin. Signals were amplified (Multiclamp 700B, Molecular Devices), low-pass filtered (1 kHz), digitized (10 kHz, Digidata 1322) and recorded (pClamp 9.2) for offline analysis.

### Fibre photometry recording

Fibre photometry was used to record calcium transients from CRH neurons in the PVN of freely moving mice. After the recovery period, animals were handled for 5 min a day for 3 days and then habituated to the optic fibre in their home cage (5 min a day) for 3 days. We recorded 20 min of CRH^PVN^ neuron activity in the homecage immediately before and for 5 min after each test to improve bleaching correction. Doric fibre photometry system consisting of two excitation LEDs (465 nm and 405 nm from Doric) controlled by an LED driver and console, running Doric Studio software (Doric Lenses). The LEDs were modulated at 208.616 Hz (465 nm) and 572.205 Hz (405 nm), and the resulting signal was demodulated using lock-in amplification. Both LEDs were connected to a Doric Mini Cube filter set (FMC6_IE(400-410)_E1(460-490)_F1(500-540)_S)_E2(555-570)_F2(580-680)_S and the excitation light was directed to the animal via a mono fibre optic patch cord (DORIC MFP_400/430/1100-0.48_2m_FC/MF2.5). The LED power was adjusted to 30 µW at the end of the patch cord. The resulting signal was detected with a photoreceiver (Newport model 2151).

### Fibre photometry data analysis

Fluorescent signal data were acquired at a sampling rate of about 100Hz (Doric system). The data were then exported to MATLAB (MathWorks) for offline analysis using custom-written scripts (https://github.com/leomol/stimbox). Briefly, raw fluorescence traces were resampled to 20 Hz to reduce data size while retaining calcium signal dynamics. A 0.5 Hz low-pass filter was then applied to suppress high-frequency noise. The 465 nm and 405 nm data were first fit individually with a second-order polynomial, and the resulting fits were subtracted to remove bleaching artifacts. Next, a least-squares linear fit was applied to the 405 nm to align it with the 465 nm channel, and then the change in fluorescence (Δ*F*) was calculated by subtracting the 405 nm Ca^2+^ independent baseline signal from the 465 nm Ca^2+^ dependent signal at each time point. The output of these transformations yielded a motion-corrected dataset of changes in GCaMP6f fluorescence. For analysis, z-score calculation was performed using the following equation: ΔF/F = (F-F_0_)/ σF, where F is the test signal, F_0_ is the median signal of the baseline, and σF is the median absolute deviation of the signal.

### Optogenetics

Following the recovery period, mice were handled for 5 min on 3 consecutive days. Then, they were habituated to the fibre-optic cable attached to the cannula (without light) for 3 additional days (15-30 min per mouse).

For in vitro recordings, a micromanipulator-attached fibre optic cable (105 μm core diameter) delivered light from a laser (Arch3.0: 593 nm, IKE-593-100-OP, IkeCool Corporation) was placed 1–2 mm away from the target area. Light intensity was calibrated by a Photodiode Power Sensor (Thorlabs). Maximally, 15 mW light was delivered to the tissue.

For in-vivo experiments, the light source (for Arch3.0: 561 nm, LRS-0561-GFO-00100-5, Laserglow Technologies) was connected to the Doric filter cube (FMC6_IE(400-410)_E1(460-490)_F1(500-540)_S)_E2(555-570)_F2(580-680)_S and the laser was directed to the animal via a mono fibre optic patch cord (DORIC MFP_400/430/1100-0.48_2m_FC/MF2.5). For Arch3.0, a yellow light (15 mW laser intensity) was used continuously for 20 seconds as the intruder was introduced to the resident’s home cage.

### Behavioural recordings and analysis

Adult male and female mice were used as residents. Resident mice encountered either a familiar or an unfamiliar intruder at each session. A familiar intruder is a conspecific from the same cage, whereas an unfamiliar intruder is from a different cage or a different strain (e.g., CD1 intruder). The test was conducted in the resident’s home cage. For video acquisition, a top-down camera (20 fps, Doric Studio) recorded all sessions under ~30–50 lux of ambient light and was synchronized to the fibre photometry recording. These videos were later analyzed using the Behavioural Observation Research Interactive Software (BORIS, version 9), and the following behaviours were scored: anogenital sniffing (directing the snout toward the anogenital area of conspecific); head sniffing (directing the snout toward the head of conspecific); torso sniffing (directing the snout toward the torso of conspecific); tail sniffing (directing the snout toward the tail of conspecific), digging (use of forepaws and sometimes hind limbs and snout to displace bedding), rearing (standing upright on hind legs with forepaws off the ground.) and grooming (repetitive licking of the body and wiping the face and head with the forepaws). Behaviour during 3 min after the introduction of the intruder was quantified and plotted. Quantification and analysis of behavioural ethograms, sliding time windows, cumulative sums of behavioural events, and transition probabilities were carried out using custom-written Python scripts. Behavioural sequences were converted into transition probability matrices by counting state-to-state transitions between consecutive behavioural events and row-normalizing counts to obtain conditional probabilities. Individual matrices were averaged across animals to generate group-level transition structures for each condition.

### Statistics

Where quantification was made, data are represented as mean ± standard error of the mean (s.e.m.). Statistical analysis was performed using GraphPad Prism 10.0. For comparisons, either two-tailed t-tests, ordinary or repeated-measures one-way ANOVA were applied as appropriate. When comparisons involved more than two groups and/or repeated factors, a two-way ANOVA was conducted, followed by Tukey’s, Sidak’s, or Holm-Sidak’s post hoc multiple-comparisons and planned comparisons test. Differences between conditions were assessed using a nonparametric permutation framework. A global test was performed using the Frobenius norm of the difference between condition-averaged matrices. Statistical significance was determined via Monte Carlo permutation (10,000 iterations), in which animal labels were randomly reassigned while preserving group sizes. P-values were computed as the proportion of permuted statistics exceeding the observed value. To identify specific transitions contributing to global differences, edge-wise comparisons were conducted using two-sided Mann–Whitney U tests on per-animal transition probabilities, with p-values corrected for multiple comparisons using the Benjamini–Hochberg false discovery rate procedure (α = 0.05). A significance threshold of p < 0.05 was applied for all analyses.

## ACKNOWLEDGEMENTS

We thank Mrs. Dinara Baimoukhametova, Ms. Alexis Passmore, Mr. Leonardo Molina, Dr. Jianjun Sun, Mrs. Cheryl Breiteneder, and Mr. Rodney Barasi for expert technical support. We also thank Dr. Matthew Hill for providing us with access to materials. We are grateful for the support of the Cumming School of Medicine Optogenetics Core Facility and the Hotchkiss Brain Institute Advanced Microscopy Platform. This work was supported by an operating grant to J.S.B. from the Canadian Institutes for Health Research and the Brain Canada Neurophotonics Platform. I.A. was supported by the Canadian Institutes for Health Research-REDI-Early career transition award.

## AUTHOR CONTRIBUTIONS

I.A. and M.P. conducted experiments, analyzed data, and prepared figures. T.S designed experiments and contributed to the paper; T.F. designed experiments, discussed results, and prepared figures. I.A. and J.S.B. designed experiments, discussed results, wrote the paper, and supervised the project.

## COMPETING INTERESTS

The authors declare they have no competing interests

## Notes

### Competing Interest Statement

The authors have declared no competing interest.

## References

1. Slavich, G.M., Roos, L.G., Mengelkoch, S., Webb, C.A., Shattuck, E.C., Moriarity, D.P., and Alley, J.C. (2023). Social Safety Theory: Conceptual foundation, underlying mechanisms, and future directions. Health Psychol. Rev. 17, 5–59. 10.1080/17437199.2023.2171900.

2. Willis, J., and Todorov, A. (2006). First Impressions Making Up Your Mind After a 100-Ms Exposure to a Face. Psychol. Sci. 17, 592–598.

3. Šabanović, M., Liu, H., Mlambo, V., Aqel, H., and Chaudhury, D. (2020). What it takes to be at the top: The interrelationship between chronic social stress and social dominance. Brain Behav. 10. 10.1002/brb3.1896.

4. Krach, S., Paulus, F.M., Bodden, M., and Kircher, T. (2010). The rewarding nature of social interactions. Preprint, 10.3389/fnbeh.2010.00022 https://doi.org/10.3389/fnbeh.2010.00022.

5. Chen, P., and Hong, W. (2018). Neural Circuit Mechanisms of Social Behavior. Preprint at Cell Press, 10.1016/j.neuron.2018.02.026 https://doi.org/10.1016/j.neuron.2018.02.026.

6. Lee, B., Ciciurkaite, G., Peng, S., Mitchell, C., and Perry, B.L. (2026). Negative social ties as emerging risk factors for accelerated aging, inflammation, and multimorbidity. Proc. Natl. Acad. Sci. U. S. A. 123. 10.1073/pnas.2515331123.

7. Tamir, D.I., and Hughes, B.L. (2018). Social Rewards: From Basic Social Building Blocks to Complex Social Behavior. Perspectives on Psychological Science 13, 700–717. 10.1177/1745691618776263.

8. Trezza, V., Campolongo, P., and Vanderschuren, L.J.M.J. (2011). Evaluating the rewarding nature of social interactions in laboratory animals. Preprint, 10.1016/j.dcn.2011.05.007 https://doi.org/10.1016/j.dcn.2011.05.007.

9. FeldmanHall, O., and Shenhav, A. (2019). Resolving uncertainty in a social world. Preprint at Nature Publishing Group, 10.1038/s41562-019-0590-x https://doi.org/10.1038/s41562-019-0590-x.

10. Kappes, A., Nussberger, A.M., Siegel, J.Z., Rutledge, R.B., and Crockett, M.J. (2019). Social uncertainty is heterogeneous and sometimes valuable. Preprint at Nature Research, 10.1038/s41562-019-0662-y https://doi.org/10.1038/s41562-019-0662-y.

11. Boucher, E.M., and Jacobson, J.A. (2012). Causal uncertainty during initial interactions. Eur. J. Soc. Psychol. 42, 652–663. 10.1002/ejsp.1876.

12. Woody, E.Z., and Szechtman, H. (2011). Adaptation to potential threat: The evolution, neurobiology, and psychopathology of the security motivation system. Preprint, 10.1016/j.neubiorev.2010.08.003 https://doi.org/10.1016/j.neubiorev.2010.08.003.

13. Hinds, A.L., Woody, E.Z., Drandic, A., Schmidt, L.A., Van Ameringen, M., Coroneos, M., and Szechtman, H. (2010). The psychology of potential threat: Properties of the security motivation system. Biol. Psychol. 85, 331–337. 10.1016/j.biopsycho.2010.08.003.

14. Füzesi, T., Daviu, N., Wamsteeker Cusulin, J.I., Bonin, R.P., and Bains, J.S. (2016). Hypothalamic CRH neurons orchestrate complex behaviours after stress. Nat. Commun. 7. 10.1038/ncomms11937.

15. Daviu, N., Füzesi, T., Rosenegger, D.G., Rasiah, N.P., Sterley, T.L., Peringod, G., and Bains, J.S. (2020). Paraventricular nucleus CRH neurons encode stress controllability and regulate defensive behavior selection. Nat. Neurosci. 23, 398–410. 10.1038/s41593-020-0591-0.

16. Rasiah, N.P., Loewen, S.P., and Bains, J.S. (2023). WINDOWS INTO STRESS: A GLIMPSE AT EMERGING ROLES FOR CRHPVN NEURONS. Preprint at American Physiological Society, 10.1152/PHYSREV.00056.2021 https://doi.org/10.1152/PHYSREV.00056.2021.

17. Füzesi, T., Rasiah, N.P., Rosenegger, D.G., Rojas-Carvajal, M., Chomiak, T., Daviu, N., Molina, L.A., Simone, K., Sterley, T.L., Nicola, W., et al. (2023). Hypothalamic CRH neurons represent physiological memory of positive and negative experience. Nat. Commun. 14. 10.1038/s41467-023-44163-5.

18. Iremonger, K.J., Power, E.M., Iremonger, K.J., and Power J Physiol, E.M. (2025). The paraventricular nucleus of the hypothalamus: a key node in the control of behavioural states. J. Physiol. 603, 2231–2243. 10.1113/JP288366#support-information-section.

19. Sterley, T.L., Baimoukhametova, D., Füzesi, T., Zurek, A.A., Daviu, N., Rasiah, N.P., Rosenegger, D., and Bains, J.S. (2018). Social transmission and buffering of synaptic changes after stress. Nat. Neurosci. 21, 393–403. 10.1038/s41593-017-0044-6.

20. Miller, C.W., Fletcher, R.J., and Gillespie, S.R. (2013). Conspecific and Heterospecific Cues Override Resource Quality to Influence Offspring Production. PLoS One 8. 10.1371/journal.pone.0070268.

21. Dawson, M., Terstege, D.J., Jamani, N., Tsutsui, M., Pavlov, D., Bugescu, R., Epp, J.R., Leinninger, G.M., and Sargin, D. (2023). Hypocretin/orexin neurons encode social discrimination and exhibit a sex-dependent necessity for social interaction. Cell Rep. 42. 10.1016/j.celrep.2023.112815.

22. Okuyama, T., Kitamura, T., Roy, D.S., Itohara, S., and Tonegawa, S. Ventral CA1 neurons store social memory.

23. Wesson, D.W., Donahou, T.N., Johnson, M.O., and Wachowiak, M. (2008). Sniffing behavior of mice during performance in odor-guided tasks. Chem. Senses 33, 581–596. 10.1093/chemse/bjn029.

24. Wachowiak, M. (2011). All in a Sniff: Olfaction as a Model for Active Sensing. Preprint, 10.1016/j.neuron.2011.08.030 https://doi.org/10.1016/j.neuron.2011.08.030.

25. Wills, G.D., Wesley, A.L., Moore, F.R., Sisemore, D.A., Wesley, A.L., Moore, F.R., and Sisemore, D.A. (1983). Social Interactions Among Rodent Conspecifics: A Review of Experimental Paradigms.

26. Tanaka, S., Young, J.W., Halberstadt, A.L., Masten, V.L., and Geyer, M.A. (2012). Four factors underlying mouse behavior in an open field. Behavioural Brain Research 233, 55–61. 10.1016/j.bbr.2012.04.045.

27. Latham, N., and Mason, G. (2004). From house mouse to mouse house: The behavioural biology of free-living Mus musculus and its implications in the laboratory. In Applied Animal Behaviour Science, pp. 261–289. 10.1016/j.applanim.2004.02.006.

28. Lever Colin, B.S.O.J. (2006). Rearing on hind legs, environmental novelty, and the hippocampal formation. Reviews in Neurosciences 17, 111–133.

29. Bels, V.L., Pallandre, J.P., Pelle, E., and Kirchhoff, F. (2022). Studies of the Behavioral Sequences: The Neuroethological Morphology Concept Crossing Ethology and Functional Morphology. Preprint at MDPI, 10.3390/ani12111336 https://doi.org/10.3390/ani12111336.

30. Donaldson, T.N., Barto, D., Bird, C.W., Magcalas, C.M., Rodriguez, C.I., Fink, B.C., and Hamilton, D.A. (2018). Social order: Using the sequential structure of social interaction to discriminate abnormal social behavior in the rat. Learn. Motiv. 61, 41–51. 10.1016/j.lmot.2017.03.003.

31. Rusu, A., Chevalier, C., de Chaumont, F., Nalesso, V., Brault, V., Hérault, Y., and Ey, E. (2023). Day-to-day spontaneous social behaviours is quantitatively and qualitatively affected in a 16p11.2 deletion mouse model. Front. Behav. Neurosci. 17. 10.3389/fnbeh.2023.1294558.

32. Zou, H., Zhang, C., Xie, Q., Zhang, M., Shi, J., Jin, M., and Yu, L. (2008). Low dose MK-801 reduces social investigation in mice. Pharmacol. Biochem. Behav. 90, 753–757. 10.1016/j.pbb.2008.06.002.

33. Doty, R.L. Odor-guided behavior in mammals.

34. Lee, W., Fu, J., Bouwman, N., Farago, P., and Curley, J.P. (2019). Temporal microstructure of dyadic social behavior during relationship formation in mice. PLoS One 14. 10.1371/journal.pone.0220596.

35. Wesson, D.W. (2013). Sniffing behavior communicates social hierarchy. Current Biology 23, 575– 580. 10.1016/j.cub.2013.02.012.

36. Chow, B.Y., Han, X., Dobry, A.S., Qian, X., Chuong, A.S., Li, M., Henninger, M.A., Belfort, G.M., Lin, Y., Monahan, P.E., et al. (2010). High-performance genetically targetable optical neural silencing by light-driven proton pumps. Nature 463, 98–102. 10.1038/nature08652.

37. Caria, A., and Dall’ò, G.M. (2022). Functional Neuroimaging of Human Hypothalamus in Socioemotional Behavior: A Systematic Review. Preprint at MDPI, https://doi.org/10.3390/brainsci12060707 10.3390/brainsci12060707.

38. Li, Y., Zeng, J., Zhang, J., Yue, C., Zhong, W., Liu, Z., Feng, Q., and Luo, M. (2018). Hypothalamic Circuits for Predation and Evasion. Neuron 97, 911-924.e5. https://doi.org/10.1016/j.neuron.2018.01.005.

39. Rasiah, N.P., Loewen, S.P., and Bains, J.S. (2023). Windows Into Stress: A Glimpse At Emerging Roles for CRH-PVN Neurons. Physiol. Rev. 103, 1667–1691. 10.1152/PHYSREV.00056.2021.

40. Denver, R.J. (2009). Structural and functional evolution of vertebrate neuroendocrine stress systems. In Annals of the New York Academy of Sciences, pp. 1–16. 10.1111/j.1749-6632.2009.04433.x.

41. Wamsteeker Cusulin, J.I., Füzesi, T., Watts, A.G., and Bains, J.S. (2013). Characterization of Corticotropin-Releasing Hormone neurons in the Paraventricular Nucleus of the Hypothalamus of Crh-IRES-Cre Mutant Mice. PLoS One 8, 1–10. 10.1371/journal.pone.0064943.

42. Ulrich-Lai, Y.M., and Herman, J.P. (2009). Neural regulation of endocrine and autonomic stress responses. Nat. Rev. Neurosci. 10, 397–409. 10.1038/nrn2647.

43. Zimmerman, C.A., Lin, Y.C., Leib, D.E., Guo, L., Huey, E.L., Daly, G.E., Chen, Y., and Knight, Z.A. (2016). Thirst neurons anticipate the homeostatic consequences of eating and drinking. Nature 537, 680–684. 10.1038/nature18950.

44. Chen, Y., Lin, Y.C., Kuo, T.W., and Knight, Z.A. (2015). Sensory Detection of Food Rapidly Modulates Arcuate Feeding Circuits. Cell 160, 829–841. 10.1016/j.cell.2015.01.033.

45. Betley, J.N., Xu, S., Cao, Z.F.H., Gong, R., Magnus, C.J., Yu, Y., and Sternson, S.M. (2015). Neurons for hunger and thirst transmit a negative-valence teaching signal. Nature 521, 180–185. 10.1038/nature14416.

46. Tirindelli, R., Dibattista, M., Pifferi, S., and Menini, A. (2009). From Pheromones to Behavior. 10.1152/physrev.00037.2008.-In.

47. Kiyokawa, Y., Kikusui, T., Takeuchi, Y., and Mori, Y. (2004). Alarm pheromones with different functions are released from different regions of the body surface of male rats. Chem. Senses 29, 35–40. 10.1093/chemse/bjh004.

48. Sterley, T.L., and Bains, J.S. (2021). Social communication of affective states. Curr. Opin. Neurobiol. 68, 44–51. 10.1016/j.conb.2020.12.007.

49. Sterley, T., Baimoukhametova, D., Füzesi, T., Zurek, A.A., Daviu, N., Rasiah, N.P., Rosenegger, D., and Bains, J.S. (2017). Social transmission and buffering of synaptic changes after stress. 10.1038/s41593-017-0044-6.

50. Anderson, D.J. (2016). Circuit modules linking internal states and social behaviour in flies and mice. Preprint at Nature Publishing Group, 10.1038/nrn.2016.125 https://doi.org/10.1038/nrn.2016.125.

51. Rho, J.-H., and Swanson, L.W. (1987). Neuroendocrine CRF motoneurons: intrahypothalamic axon terminals shown with a new retrograde-Lucifer-immuno method.

52. Rajamanickam, S., and Justice, N.J. (2022). Hypothalamic corticotropin-releasing factor neurons modulate behavior, endocrine, and autonomic stress responses via direct synaptic projections. Curr. Opin. Endocr. Metab. Res. 26. 10.1016/j.coemr.2022.100400.

53. Füzesi, T., Daviu, N., Wamsteeker Cusulin, J.I., Bonin, R.P., and Bains, J.S. (2016). Hypothalamic CRH neurons orchestrate complex behaviours after stress. Nat. Commun. 7, 1–14. 10.1038/ncomms11937.

54. Hunt, A.J., Dasgupta, R., Rajamanickam, S., Jiang, Z., Beierlein, M., Chan, C.S., and Justice, N.J. (2018). Paraventricular hypothalamic and amygdalar CRF neurons synapse in the external globus pallidus. Brain Struct. Funct. 223, 2685–2698. 10.1007/s00429-018-1652-y.

55. Haller, J., Barna, I., and Baranyi, M. (1995). Hormonal and metabolic responses during psychosocial stimulation in aggressive and nonaggressive rats. Psychoneuroendocrinology 20, 65– 74.

56. Xu, S., Yang, H., Menon, V., Lemire, A.L., Wang, L., Henry, F.E., Turaga, S.C., and Sternson, S.M. (2020). Behavioral state coding by molecularly defined paraventricular hypothalamic cell type ensembles. Science (1979). 370. 10.1126/science.abb2494.

57. Zimmerman, C.A., Lin, Y.C., Leib, D.E., Guo, L., Huey, E.L., Daly, G.E., Chen, Y., and Knight, Z.A. (2016). Thirst neurons anticipate the homeostatic consequences of eating and drinking. Nature 537, 680–684. 10.1038/nature18950.

58. Chen, Y., Lin, Y.C., Kuo, T.W., and Knight, Z.A. (2015). Sensory Detection of Food Rapidly Modulates Arcuate Feeding Circuits. Cell 160, 829–841. 10.1016/j.cell.2015.01.033.

59. Kim, J., Lee, S., Fang, Y.Y., Shin, A., Park, S., Hashikawa, K., Bhat, S., Kim, D., Sohn, J.W., Lin, D., et al. (2019). Rapid, biphasic CRF neuronal responses encode positive and negative valence. Nat. Neurosci. 22, 576–585. 10.1038/s41593-019-0342-2.

